# A novel stimulus paradigm for simultaneous recording of monaural and binaural frequency following response for identification of binaural interaction component

**DOI:** 10.1101/2020.03.09.983205

**Authors:** Srividya Grama Bhagavan, Mohan Kumar Kalaiah

## Abstract

The objective of the study was to investigate whether monaural frequency following response (FFR) of right and left ear and binaural FFR could be obtained in the same recording using a novel stimulus presentation paradigm, for the purpose of identification the BIC. Twenty six young adults participated in the study. The FFR was recorded for 220 Hz pure-tone using a novel stimulus paradigm. The pure-tone was presented sequentially to two ears. Initially, the pure-tone was presented to the right ear, then to both ears, and finally to the left ear. The FFR could be elicited from all participants (all three responses: right ear, left ear, and both ears) in the same recording using the novel stimulus presentation paradigm used in the present study. The novel stimulus presentation paradigm used in the present study could be used for obtaining monaural and binaural FFRs in the same recording for identification of BIC.

## Introduction

The frequency-following response (FFR) is an auditory evoked potential which follows the periodicity of the stimuli (Moushegian et al, 1973). It reflects the synchronous activity of axonal and dendritic potentials generated by neurons in the lateral lemniscus and inferior colliculus of the brainstem (Smith et al, 1975; Møller, 1998). Recently, the FFR has gained popularity, and it is being used to gain insight into abnormal encoding of complex sounds in clinical population (Cunningham et al, 2001; Rocha-Muniz et al, 2012; Billiet & Bellis, 2011; Hornickel & Kraus, 2013), effects of ageing on encoding of sound at sub-cortical structures (Anderson et al, 2012; Bidelman et al, 2014; Parthasarathy & Bartlett, 2012), binaural processing abilities (Clinard et al, 2017; Graydon et al, 2019; Wilson & Krishnan, 2005), individual differences in speech understanding and neurobiology of music (Bidelman, 2013; Bidelman et al, 2011; Bones et al, 2014), and neuroplasticity of learning and language experience (Bidelman & Alain, 2015).

Studies aimed to investigate binaural processing at subcortical level have commonly measured masking level difference (MLD) or binaural interaction component (BIC) using the FFR (Clinard et al, 2017; Graydon et al, 2019; Wilson & Krishnan, 2005). Traditionally, the MLD is measured by computing the difference in hearing thresholds between homophasic and antiphasic conditions. In order to identify the BIC, typically auditory evoked potentials are recorded from right ear, left ear, and both ears. Finally, the BIC is measured as the difference between the sum of the right and left ear responses and the binaural response (Dobie & Berlin, 1979). Studies have shown that, the BIC can be recorded using several auditory evoked potentials such as auditory brainstem response (Gopal & Pierel, 1999; McPherson & Starr, 1993; Delb et al, 2003), middle latency response (Abdollahi et al, 2017; Kelly-Ballweber & Dobie, 1984; McPherson & Starr, 1993), late latency response (McPherson & Starr, 1993), and FFR (Krishnan & McDaniel, 1998; Uppunda et al, 2015).

Earlier studies which investigated the BIC using electrophysiological measures have traditionally recorded responses from right ear, left ear, and both ears in separate recordings (Dobie & Berlin, 1979; McPherson & Starr, 1993; Van Yper et al, 2015; Wrege & Starr, 1981). Although the traditional method is suitable for identifying the BIC, it has few limitations. Firstly, the level of residual noise in the averaged waveform could be different across recordings. In addition, since responses from right ear, left ear, and both ears are recorded separately it would increase the test time. Therefore, the aim of present study was to record all three responses in the same recording using a novel stimulus presentation paradigm. The advantage of novel stimulus presentation paradigm is that a single recording would be sufficient to obtain response of right ear, left ear, and both ears, for the purpose of identification the BIC. In the present study, the FFR was recorded using the novel stimulus presentation paradigm. The FFR was selected as it is gaining popularity in recent years as an objective tool for understanding binaural processing at subcortical levels.

## Method

### Participants

Twenty-six young adults (female=19, male=7) aged between 20 to 40 years (mean age= 25.15 years) participated in the study. Air conduction pure tone threshold of all participants was less than 15 dB HL at octave frequencies between 250 Hz and 8000 Hz in both ears. Immittance evaluation showed normal findings in both ears, i.e., ‘A’ type tympanogram with acoustic reflex present at normal levels. None of the participants had history of otological or neurological problems, exposure to loud noise, ototoxic medication, and metabolic disorders such as diabetes and hypertension. The study was approved by the institutional ethics committee of Kasturba Medical College, Mangalore. Informed consent was obtained from all participants before their participation in the study.

### Stimuli

Pure tones of 220 Hz with a duration of 50 msec and 100 msec were used for the recording of the frequency following response. Pure tones were generated using MATLAB at a sampling rate of 44100 Hz and 16-bit resolution. Onset and offset of pure tones were windowed (using 10 msec Hamming window) to reduce spectral splatter. Waveforms of stimuli used to elicit the FFR are shown in figure 1. Two methods of stimulus presentation were used to elicit the FFR, referred to as ‘stimulus without gap’ and ‘stimulus with gap’ conditions. In ‘stimulus without gap’ condition, two pure tones with a duration of 100 msec were used to elicit the frequency following response. Pure tones were presented sequentially in two ears with partial overlap of pure tones (50 msec). The pure tone was always presented first to the right ear, then to the left ear after a lag of 50 msec. Thus, in the initial 50 msec, the pure tone was heard only in the right ear, following this pure tones were heard in both ears for 50 msec, and finally, the pure tone was heard only in the left ear for 50 msec. In ‘stimulus with gap’ condition, pure tones with a duration of 50 msec were used to elicit the FFR. Initially, the pure tone was presented only in the right ear, after a silence period of 10 msec pure tones were presented to both ears simultaneously, and finally, the pure tone was presented only in the left ear after a silence period of 10 msec. In addition, the onset phase of the pure tone presented to the left ear was varied in binaural stimulation condition. The onset phase of the pure tone in the left ear was 0 and 180 degree, referred to as ‘same-phase’ condition and ‘opposite-phase’ condition, respectively.

**Figure 1.**
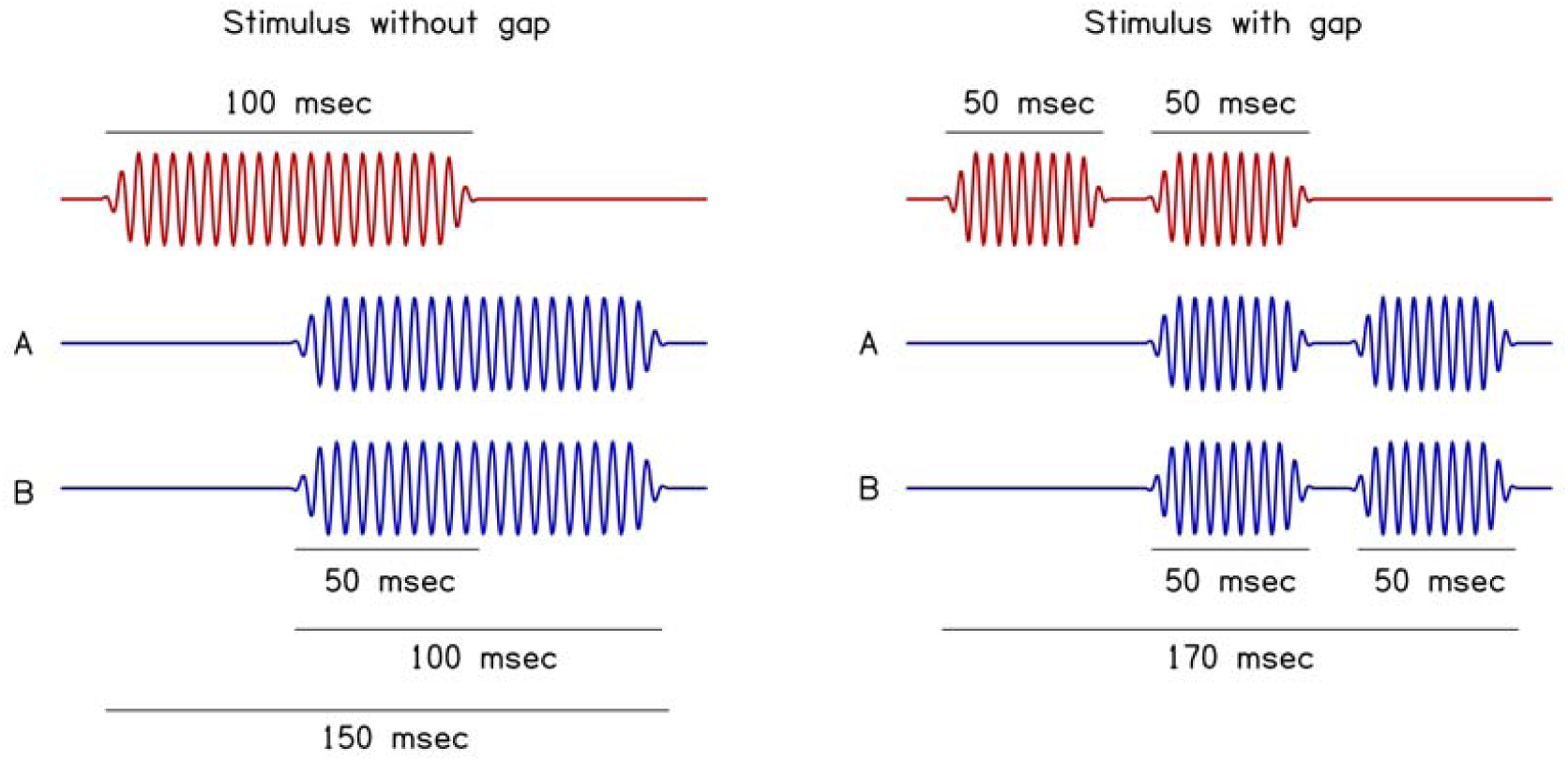
Waveforms of stimuli used for eliciting the FFR. The left panel shows waveforms of stimuli used in ‘stimulus without gap’ condition and the right panel shows waveforms of stimuli used in ‘stimulus with gap’ condition. The red coloured waveform in both conditions represent pure tone presented to the right ear, and blue coloured waveforms represent stimuli presented to the left ear. Waveform labelled ‘A’ has same phase compared to the right ear and waveform labelled ‘B’ has opposite phase compared to the right ear.

### Recording of the frequency following response

The FFR was recorded using IHS Smart EP-CAM evoked potential system. During the recording of the FFR, participants were made to sit comfortably on a reclining chair, and they were instructed to relax and minimise extraneous body movements to reduce unwanted artifacts. The EEG was differentially recorded from the scalp using gold-coated disc electrodes. The non-inverting electrode was placed on vertex, inverting electrode on the nape of the neck, and ground electrode was placed on low forehead. The absolute electrode impedance and inter-electrode impedance was maintained below 5 kOhm and 2 kOhm, respectively. Pure tones were presented at 80 dB SPL to both ears of participants using Ear-Tone ER-3A insert phones. The ongoing electrical activity (EEG) was band-pass filtered (60 Hz to 3000 Hz) and digitised at a sampling rate of 10,000 Hz and stored for off-line analysis. During offline analysis, the continuous EEG was band-pass filtered between 80 Hz and 1500 Hz using finite impulse response filter (slope=24 dB/octave). The filtered EEG was segmented to obtain epochs of 2300 or 2700 msec, including a pre-stimulus baseline of 60 msec or 80 msec respectively. The segmented EEG was subjected to baseline correction and artifact removal. Baseline correction was performed by considering pre-stimulus activity between −20 msec and 0 msec as a baseline. FieldTrip toolbox for MATLAB (Donders Centre for Cognitive Neuroimaging in Nijmengen, Netherlands) (Oostenveld et al, 2011) was used for band-pass filtering and baseline correction of EEG during offline analysis. Finally, segments with excessive noise or amplitude variation greater than 50 µV were removed, and averaging was done to obtain averaged waveform. Stimulus and recording parameters used for eliciting the FFR are shown in table 1.

**Table 1:** Stimulus protocol used in the present study for the recording of the FFR.

| **Parameter** | **‘Stimulus without gap’**<br>**condition** | **‘Stimulus with gap’**<br>**condition** |
| --- | --- | --- |
| Stimulus | 1. 100 msec pure tone (220 Hz, onset phase = 0 degree)<br>2. 100 msec pure-tone (220 Hz, onset phase = 180 degrees) | 1. 50 msec pure tone (220 Hz, onset phase = 0 degree)<br>2. 50 msec pure-tone (220 Hz, onset phase = 180 degrees) |
| Number of sweeps | 2000 | 2000 |
| Inter-stimulus interval | 230 msec | 270 msec |
| Polarity | Single polarity | Single polarity |
| Intensity | 80 dB SPL | 80 dB SPL |
| Filters | 60 Hz to 3000 Hz | 60 Hz to 3000 Hz |
| Amplification | 1,00,000 | 1,00,000 |
| Analysis time window | 230 msec (including the pre-stimulus window of 60 msec) | 270 msec (including the pre-stimulus window of 80 msec) |
| Number of channels | One | One |

### Data analysis

The averaged waveforms obtained from all participants were subjected to spectral analysis. Spectral analysis was performed on segments of averaged waveform comprising pre-stimulus activity, the monaural response of right ear and left ear, and binaural response. Waveforms were subjected to fast fourier transformation analysis to obtain the spectrum of response waveform. From the spectrum, peak amplitude was measured in the frequency regions adjacent to the frequency of pure-tone (referred as peak F_0_ amplitude). In addition, average amplitude was computed based on amplitude of the FFR in the frequency regions adjacent to the frequency of pure-tone (referred as average F_0_ amplitude). In addition, the root-mean-square amplitude was computed for segments of averaged waveform comprising pre-stimulus activity, the monaural response of right and left ears, and binaural response.

## Results

The FFR could be elicited from all participants using both methods of stimulus presentation. Figure 2 shows averaged waveforms of the FFR obtained from one participant using both methods of stimulus presentation. The FFR elicited during monaural and binaural stimulation were different in both methods of stimulus presentation. In general, for both methods of stimulus presentation, the amplitude of binaural FFR was greater than monaural FFR in ‘same-phase’ condition. Whereas, in ‘opposite-phase’ condition, the amplitude of binaural FFR was comparable to monaural FFR, while, the waveform of binaural FFR in ‘opposite-phase’ condition was distinct than monaural FFR. Figure 3 shows mean RMS amplitude, average F_0_ amplitude, and peak F_0_ amplitude of monaural FFR (right ear and left ear) and binaural FFR for both methods of stimulus presentation. In ‘stimulus with gap’ condition, the mean RMS amplitude, average F_0_ amplitude, and peak F_0_ amplitude of monaural FFR were similar in both ears. In contrast, the mean RMS amplitude, average F_0_ amplitude, and peak F_0_ amplitude were higher in the right ear compared to left ear in ‘stimulus without gap’ condition. Further, in the right ear, the mean RMS amplitude, average F_0_ amplitude, and peak F_0_ amplitude were similar for both methods of stimulus presentation.

**Figure 2.**
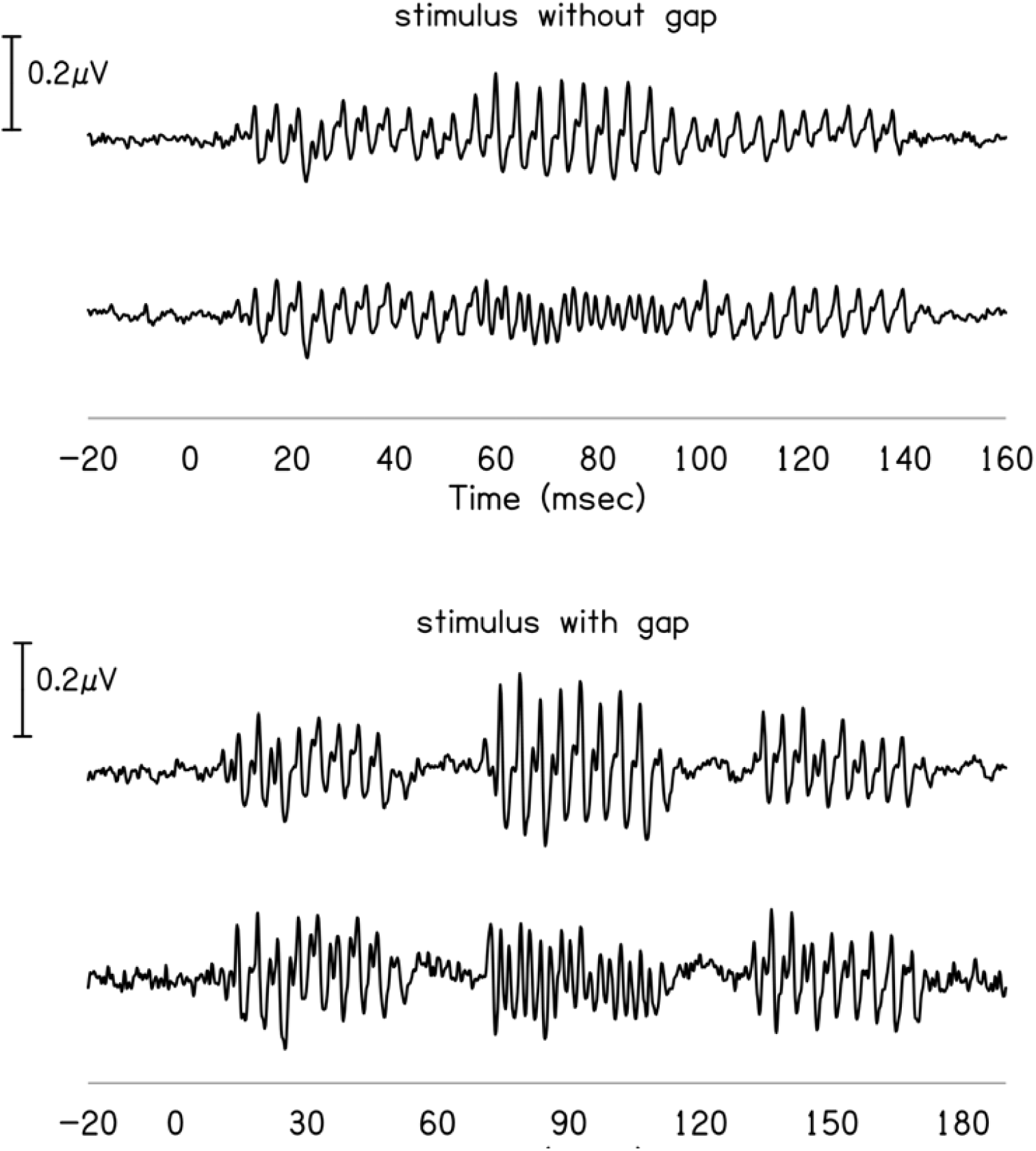
Waveforms of the binaural FFR elicited from one participant using two methods of stimulus presentation. Waveforms labelled ‘A’ and ‘B’ in the top panel represents response obtained with ‘same phase’ and ‘opposite phase’ respectively.

**Figure 3.**
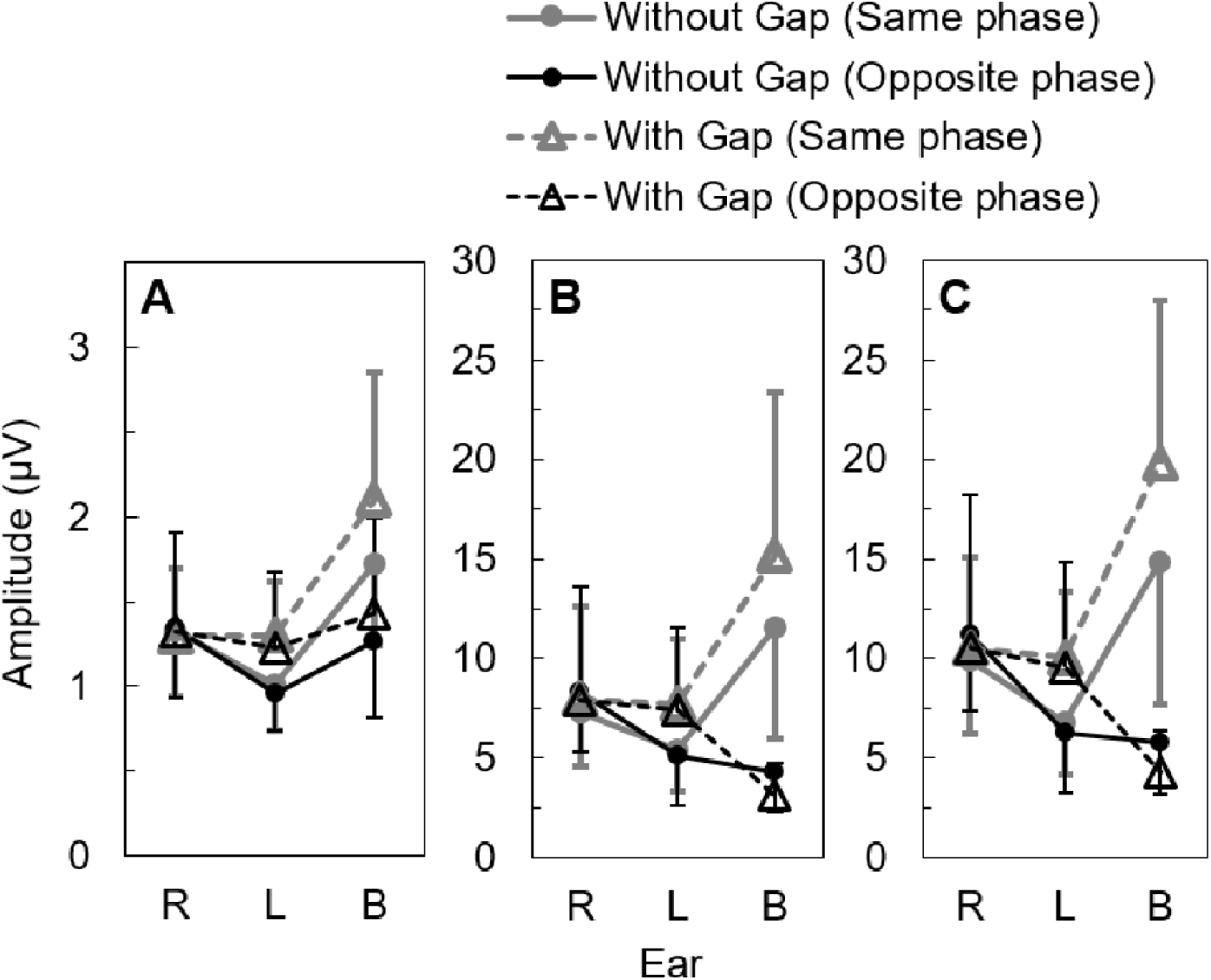
Mean amplitude of the response for the right ear, left ear, and both ears for two methods of stimulus presentation. Panel A shows the mean root mean square amplitude of the response. Panel B and Panel C shows the mean average F_0_ amplitude and mean peak F_0_ amplitude of the response.

To investigate whether the amplitudes of right and left ears are significantly different between the two methods of stimulus presentation, data was subjected to further statistical analysis. Initially, paired-samples t-test was carried out separately for right and left ears, to investigate whether the response amplitudes are significantly different between the ‘same phase’ and ‘opposite phase’ conditions. It showed no significant difference for the RMS amplitude, average F_0_ amplitude, and peak F_0_ amplitude between two conditions for both right and left ears. Since amplitudes were not significantly different between ‘same phase’ and ‘opposite phase’ conditions, these data were combined and subjected to further analysis. To investigate whether mean amplitudes are significantly different between two ears, paired-sample t-test was carried out separately for two methods of stimulus presentation. It revealed no significant difference for the mean RMS amplitude [t(25)=0.719, p=0.479], average F_0_ amplitude [t(25)=0.517, p=0.610], and peak F_0_ amplitude [t(25)=0.670, p=0.509] between two ears in ‘stimulus with gap’ condition. In contrast, in ‘stimulus without gap’ condition, the mean amplitudes of response were reduced significantly in the left ear compared to the right ear (RMS amplitude [t(25)=7.289, p<0.001]; F_0_ amplitude [t(25)=4.649, p<0.001]; peak F_0_ amplitude [t(25)=5.067, p<0.001]).

## Discussion

In the present study, the FFR could be recorded from all participants using both methods of stimulus presentation. Further, both binaural FFR and monaural FFR of both ears could be recorded in the same recording from all participants. This finding in the present study showed that by using the stimulus paradigm used in the present study, the FFR could be recorded from both ears in the same recording. Simultaneous recording of monaural and binaural FFR could be beneficial for assessing binaural processing at the neural level (e.g., binaural interaction component). In consonance with findings of the present study, few investigations have demonstrated that auditory evoked potentials can be recorded from both ears simultaneously in the same recording using suitable stimulus presentation paradigms (Maruthy et al, 2019; John et al, 1998; Ishida & Stapells, 2012). Recently, Maruthy et al. (2019) recorded click ABR using a novel stimulus paradigm and showed that click ABR could be recorded from both ears simultaneously in the same recording. Further, results showed that the click ABR of both ears obtained using novel stimulus paradigm was comparable to click ABR elicited using the conventional method. Ishida and Stapells (2012) recorded 40 Hz and 80 Hz ASSR using multiple stimuli presented simultaneously to one ear or both ears and showed that ASSR could be recorded from both ears simultaneously in the same recording. The amplitude of 80 Hz ASSR recorded by presenting multiple stimuli to both ears was similar to multiple stimuli to one ear condition. However, the amplitude of 40 Hz ASSR with multiple stimuli to both ears was smaller than multiple stimuli to one ear. These findings show that auditory evoked potentials can be recorded from both ears simultaneously in the same recording.

The present study compared the FFR recorded using two methods of stimulus presentation (‘stimulus without gap’ and ‘stimulus with gap’). In ‘stimulus without gap’ condition, the amplitude of the FFR in the left ear was reduced compared to the right ear. On the other hand, in ‘stimulus with gap’ condition, the amplitude of the FFR was similar in both ears. The reduction in amplitude of the FFR in the left ear, in ‘stimulus without gap’ condition, could be attributed to two reasons. First, reduced amplitude in the left ear could be a consequence of the order of presentation of pure-tones to the right and left ears, i.e., order effect. In the present study, the FFR was elicited from two ears by presenting pure-tone first to the right ear and then to the left ear. The first pure-tone, which was presented to the right ear, could have elicited stronger response compared to the second pure-tone. Thus, the amplitude of the FFR in the left ear was reduced compared to the right ear. Secondly, the reduced amplitude of the FFR in the left ear could be a consequence of the reduction in the neural firing rate over time due to adaptation of neurons contributing to the generation of the FFR (Gockel et al, 2013; Gockel et al, 2015). In the present study, the response of right and left ears were obtained at time intervals between 10 msec and 60 msec and 110 msec and 160 msec post stimulus onset, respectively. Gockel et al, (2013; 2015) recorded the FFR using long duration pure tone and compared the amplitude of response at beginning, middle, and ending portions. Findings showed that the amplitude of the FFR was largest in start portion and smallest in end portion. They attributed the reduction in the amplitude of the FFR over time to the adaptation of neurons generating the response. Result of the present study was comparable to findings reported by (Gockel et al, 2013; Gockel et al, 2015), where both investigations showed greater amplitude for the FFR at earlier time intervals and a reduction in its amplitude over time over time. Therefore, reduced amplitude in the left ear could be attributed to a decrease in neural activity over time due to the adaptation of neurons. In contrast, the amplitude of the FFR in the left ear was similar to the right ear in ‘stimulus with gap’ condition. This increase in the amplitude of the FFR in the left ear could be attributed to the recovery of neurons from adaptation. In ‘stimulus with gap’ condition, a short period of silent interval was introduced between pure-tones eliciting right and left ear responses. Thus, the silent interval between pure-tones would have facilitated the recovery of neurons from adaptation leading to an increase in the amplitude of the FFR in the left ear.

The present research attempted to record monaural FFR of both ears and binaural FFR in the same recording using a novel method of stimulus presentation. Results showed that, by presenting stimuli to the right ear, left ear, and both ears sequentially both monaural and binaural frequency following response could be recorded from all participants. This method of stimulus presentation could be useful during the recording of binaural interaction component of the FFR. However, in the present study the binaural interaction component was not measured. In addition, the monaural FFR of right and left ears were not compared with conventional recording procedures.

